# The mechanics of correlated variability in segregated cortical excitatory subnetworks

**DOI:** 10.1101/2023.04.25.538323

**Authors:** Alex Negrón, Matthew P. Getz, Gregory Handy, Brent Doiron

**Author notes:** These authors contributed equally (Alex Negrón, Matthew P. Getz). These authors contributed equally (Gregory Handy, Brent Doiron).

## Abstract

Understanding the genesis of shared trial-to-trial variability in neural activity within sensory cortex is critical to uncovering the biological basis of information processing in the brain. Shared variability is often a reflection of the structure of cortical connectivity since this variability likely arises, in part, from local circuit inputs. A series of experiments from segregated networks of (excitatory) pyramidal neurons in mouse primary visual cortex challenge this view. Specifically, the across-network correlations were found to be larger than predicted given the known weak cross-network connectivity. We aim to uncover the circuit mechanisms responsible for these enhanced correlations through biologically motivated cortical circuit models. Our central finding is that coupling each excitatory subpopulation with a specific inhibitory subpopulation provides the most robust network-intrinsic solution in shaping these enhanced correlations. This result argues for the existence of excitatory-inhibitory functional assemblies in early sensory areas which mirror not just response properties but also connectivity between pyramidal cells.

## 1 Introduction

Determining a structure – function relationship in a cortical circuit is a central goal in many neuroscience research programs. While the trial averaged responses of a network to a fixed stimulus or repeated behavior does give some information about the underlying circuit, the dynamic or trial-to-trial fluctuations of neuronal activity provides another important glimpse into network structure (Urai et al., 2022). Such neuronal variability is a salient feature of cortical responses (Faisal et al., 2008), and of particular interest is how that variability is distributed over a population of neurons (Cohen and Kohn, 2011). The shared fluctuations of a pair of neurons, termed *noise correlations*, are often thought to reflect the circuit structure of the network within which the neuron pair is embedded (Doiron et al., 2016; Ocker et al., 2017). Understanding how neural variability is shaped by the connections and local circuit dynamics can provide rich insight into the structure and function of cortical circuits.

An early hope was that pairwise correlations in neuronal activity could be used to infer the underlying connectivity in a straightforward fashion (Mishchencko et al., 2011; Roudi et al., 2015). Indeed, experiments in the mouse primary visual cortex (V1) demonstrated that the magnitude of the pairwise correlation between two pyramidal cells increases with their probability of connection (Ko et al., 2011; Cossell et al., 2015). Theoretically, this result can be thoroughly explained in a weakly coupled excitatory network, where correlations are dominantly determined by direct, monosynaptic connections. However, recent experiments investigating the functional properties of two distinct subpopulations of pyramidal cells in mouse V1 complicate this narrative (Kim et al., 2018b). These subpopulations project to separate downstream higher visual areas and are inter-connected with lower probability than that of randomly sampled pyramidal cells within V1. Despite this weak connectivity, it was found that the correlations between these distinct subpopulations were much higher than predicted by their sparse inter-connectivity. In fact, the magnitude of the correlated variability across the two subpopulations approached that between any randomly chosen pair of excitatory neurons. In this same vein, another experiment examining callosal projection neurons in mouse V1 found that these cells also cluster and connect more strongly as a class (Hagihara et al., 2021). Yet their correlated variability is similar when comparing within-class and out-of-class, again illustrating that significant, positive correlations can persist in the absence of direct strong connections. In total, these results are at odds with previous intuition, namely that this anatomical segregation would correspond to a functional one as well.

Theoretical work has also highlighted how the simplistic structure-dynamics relationship originally put forth can break down. For example, it has been shown that inferring connectivity from activity becomes difficult as recurrent connection strengths grow and inhibition is required to stabilize the network (Das and Fiete, 2020; Biswas and Fitzgerald, 2022). Most notably, a densely connected network with strong synaptic weights that exists in the so-called balanced state, a robust parameter regime where excitatory and inhibitory inputs to a neuron largely cancel out, results in near-zero average correlations (Renart et al., 2010; Rosenbaum et al., 2017). Nonetheless, progress has been made in overcoming these difficulties, with studies having developed methods for linking connectivity motifs to the structure of correlations in arbitrarily large networks (Pernice et al., 2011; Trousdale et al., 2012; Ocker et al., 2017)).

In this work, we seek to apply some of these techniques to characterize the neural circuit properties which could explain the significant positive shared variability across segregated cortical subpopulations observed in Kim et al. (2018b). With the use of mean field circuit models we show that the solution depends on the dynamical regime of the circuit, and relies on the structure of inhibition. In a weakly coupled regime, correlations can be characterized through inheritance from outside sources, or increased through shared inhibitory inputs. By contrast, in a strongly coupled regime, shared inhibition would largely act to anticorrelate activity across the populations. Critically, we show that this anticorrelation can be mitigated if inhibition is similarly clustered with excitation, forming instead excitatory-inhibitory assemblies. Additionally, this regime of strongly coupled dynamics with clustered inhibition provides the most robust solution space to explain the elevated correlations. This prediction further suggests that other apparent correlation conundrums could be solved by supplementing excitatory recordings with activity from inhibitory neurons.

## 2 Results

### 2.1 Segregated synaptic wiring does not produce segregated functional responses

Our work is motivated by an apparent inconsistency in a series of experimental studies exploring the relation between the recurrent circuitry and functional responses of neuronal populations in sensory neocortex. Ko et al. (2011) and Cossell et al. (2015) used a combination of *in vivo* population imaging and *in vitro* electrophysiology to show that the activity correlations between pairs of pyramidal neurons in mouse primary visual cortex (V1) increase monotonically with the probability of there existing synaptic connections between them. Later work from the same group (Kim et al., 2018b) investigated two excitatory populations in mouse V1: neurons that are either anterolateral (AL)- or posteromedial (PM)-projecting. Despite neurons being in close spatial proximity to each other, these neuronal subpopulations exhibit high within group connectivity (prob. AL ↔ AL connection ∼ 0.21, prob. PM ↔ PM connection ∼ 0.18) and low between group connectivity (prob. AL → PM connection ∼ 0.04, prob. PM → AL connection ∼ 0.05). To streamline our presentation we will label these two populations *E*_1_ and *E*_2_ (Fig. 1A). Given the low connection probability between *E*_1_ and *E*_2_ and the established relation between connectivity and activity correlations shown in Ko et al. (2011) and Cossell et al. (2015), one would predict that the degree of correlations between the activities of *E*_1_ and *E*_2_ would be low (Fig. 1B, held out light green square; from Kim et al. (2018b) we estimate this value to be approximately -0.05). However, Kim et al. (2018b) reported substantially higher than predicted mean *E*_1_ and *E*_2_ correlations (Fig. 1B, dark green square; Kim et al. (2018b) measured it to be about 0.027, close to the within-group values of about 0.035-0.04). In total, while pyramidal neurons in mouse V1 projecting to distinct targets show segregated synaptic connectivity, the degree of functional segregation between these subpopulations is below what is expected.

**Figure 1.**
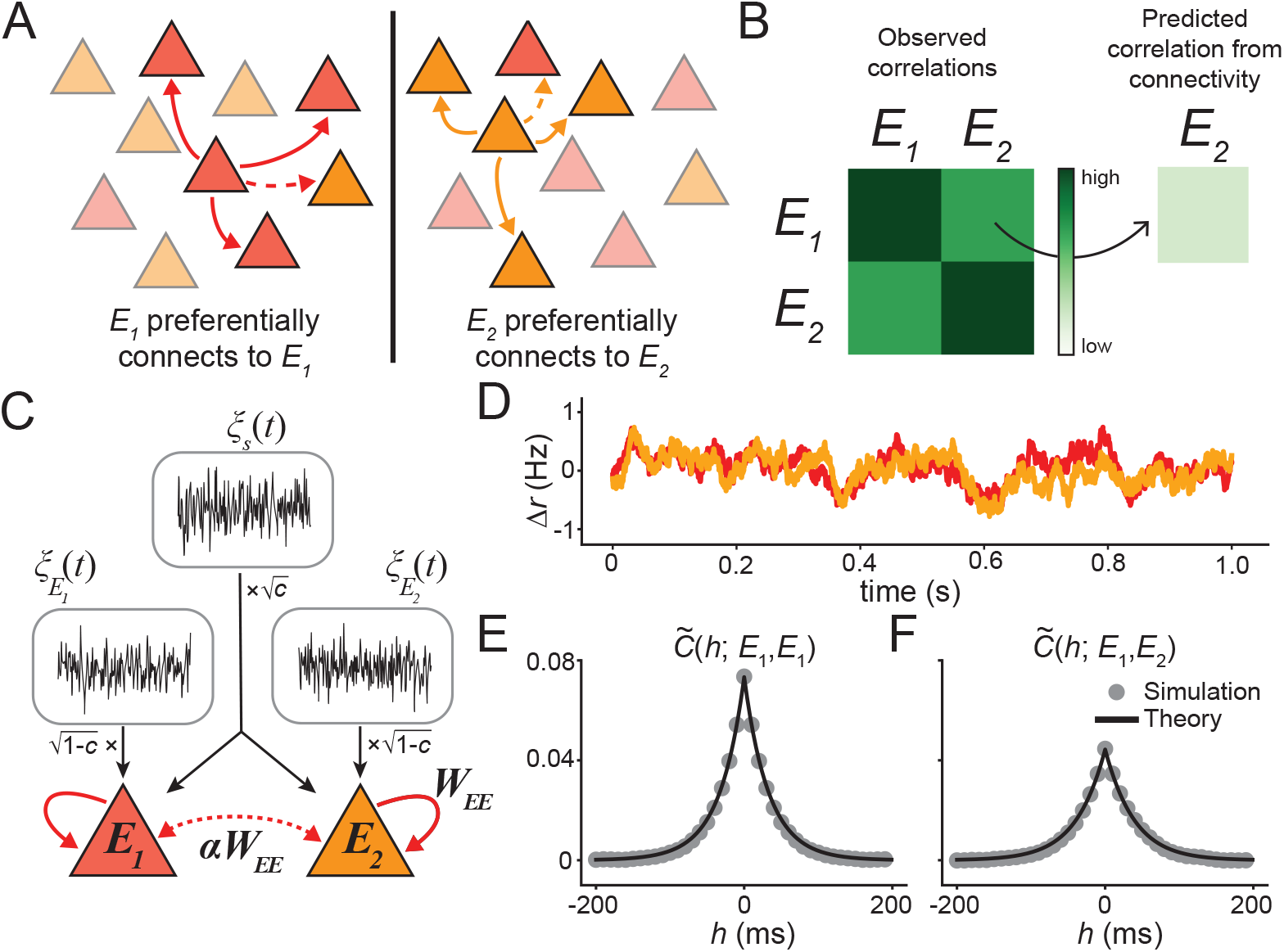
Mean field model of segregated E populations. **A:** Illustration of experimentally observed connectivity motif; the red (*E*_1_) and orange (*E*_2_) populations connect with lower probability than average. **B:** Schematic of main experimental observations: *E*_1_−*E*_2_ correlations were higher than would be predicted from their low connectivity. **C:** Model schematic. Black traces and arrows denote noise sources. Red arrows indicate excitatory recurrent connections where the dashed line connotes weakened connection strength. Feedforward stimulus drive omitted for clarity. **D:** Example realization of network activity to a sustained, fixed stimulus. Colors as in (A). **E:** *E*_1_ auto-correlation function and **F:** *E*_1_−*E*_2_ cross-correlation function for the illustrated rate traces. For panels D, E and F: *c* = 0.5.

The central goal of our study is to put forth a circuit-based model framework that can robustly and self-consistently account for both of these experimental observations. It is important to note that Kim et al. (2018b) only considered total correlations (of the raw neural activity traces) in computing this expected correlation value. However, given the similarities observed in the signal and noise correlation structure in both this and previous studies (Ko et al., 2011; Kim et al., 2018b; Hagihara et al., 2021), we focus here on noise correlations which relate more directly to the underlying structure of connectivity (Ocker et al., 2017).

### 2.2 A circuit model of fluctuations in segregated subpopulations

To study the structure of correlations in anatomically segregated networks and investigate the possible mechanisms responsible for the unexpectedly enhanced correlations, we consider a phenomenological dynamic mean field model for the aggregate activity of each neural population (Renart et al., 2004; Getz et al., 2022; Kanashiro et al., 2017). Assuming that the network has a steady state solution (**r**_ss_), the linearized dynamics of population *A* around this equilibrium are given by (see Section 5 for additional details):

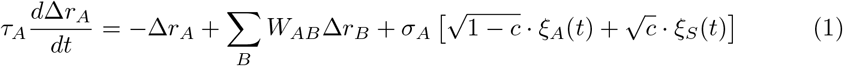

where Δ*r*_*A*_ = *r*_*A*_ − *r*_ss,*A*_, *τ*_*A*_ is a time constant, and *W*_*AB*_ is the effective strength of connections from population *B* to *A*. For the purely excitatory network *A* and *B* range over *E*_1_ and *E*_2_; when inhibitory connections are included in later sections *A* and *B* will include those as well. The stochastic processes *ξ*_*A*_(*t*) and *ξ*_*S*_(*t*) represent private and shared global fluctuations, respectively, modelling stochastic inputs that are external to the network. *ξ*_*A*_(*t*) and *ξ*_*S*_(*t*) are taken to be independent Gaussian processes with ⟨*ξ*(*t*)⟩ = 0 and ⟨*ξ*(*t*)*ξ*(*t*^*′*^)⟩ = *δ*(*t* − *t*^*′*^). The parameter *c* ∈ [0, 1] scales the proportion of shared noise relative to private noise (Fig. 1C), while *Σ*_*A*_ *>* 0 represents the total intensity of the external fluctuations given to population *A*.

We make two assumptions: 1) the network has a stable solution *r*_*ss*_ about which the population dynamics fluctuate (Fig. 1D), and 2) connections within and inputs to the network are symmetric across the two *E* populations, with 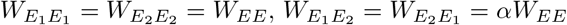, and 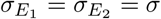. Note that parameter 0 *< α*≪1 represents the degree to which the inter-population connections are weaker than the within-population connections (Fig. 1C). Since the system of recurrently coupled stochastic differential equations in Eq. 1 is a multi-dimensional Ornstein-Uhlenbeck (OU) process, we can derive (see Section 5) an analytical formula for its stationary autocovariance function

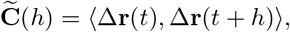

which agrees well with numerical simulations (Fig. 1E, F). Further, and of particular interest in this work, is the *long-time covariance matrix* defined as

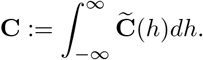

This may be expressed (see Methods 5.2) as

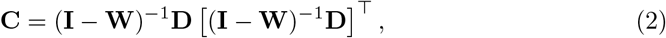

where **W** is a matrix of effective connection strengths and **D** is a matrix that scales the fluctuations. We define the correlations between *E*_1_ and *E*_2_ as

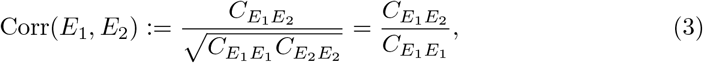

where *C*_*AB*_ is an element of **C** and the second equality follows by the assumed symmetry in the system. This framework enables us to formalize the motivating question of our study: what are the mechanisms that enable higher than expected correlations across anatomically segregated populations? For the sake of specificity, we choose the threshold Corr(*E*_1_, *E*_2_) *>* 0.6 as an approximation of the ratio of mean across-population to within-population noise correlations in Kim et al. (2018b).

### 2.3 Inheritance model of correlations between weakly coupled excitatory populations

We begin by exploring how the strength of recurrent excitation (*W*_*EE*_) and the proportion of fluctuations that are shared (*c*) shape correlations between the segregated *E* populations. In this section, to ensure that the network admits a stable activity solution we require *W*_*EE*_ *<* 1, else recurrent excitation would lead to runaway activity. Note that while we allow *W*_*EE*_ to vary, we maintain the concept of segregated populations by keeping *α* small and fixed. We find that while increasing *W*_*EE*_ leads to moderate increases in Corr(*E*_1_, *E*_2_), a much more significant increase occurs by increasing *c* (Fig. 2A).

**Figure 2.**
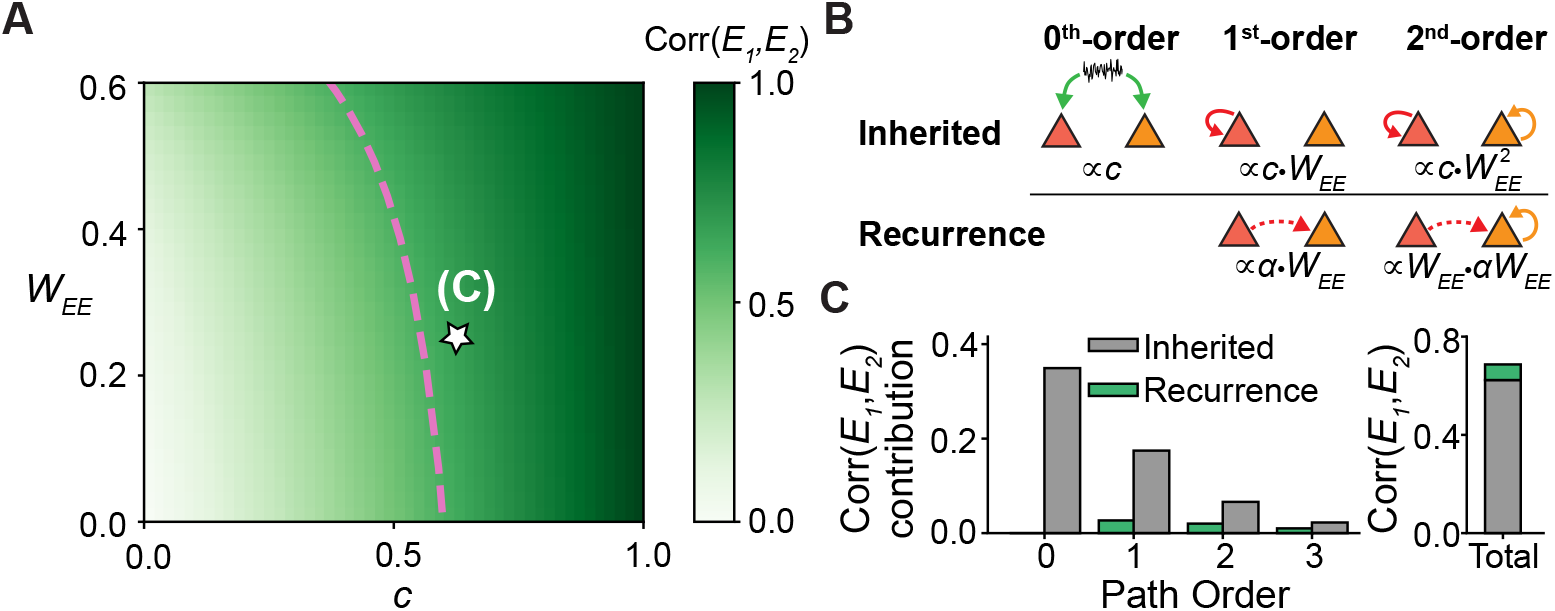
Highly correlated regime in weakly coupled excitatory network relies on correlated feedforward inputs. **A:** Corr(*E*_1_, *E*_2_) as a function of *W*_*EE*_ and the magnitude of shared input noise *c*. Dashed pink line indicates Corr(*E*_1_, *E*_2_) = 0.6, approximating the value reported in Kim et al. (2018b). **B:** Schematic of example synaptic paths through the network, along with their contribution to the cross-covariance, relating to the path expansion Eq. 4. The inherited row refers to correlated paths stemming from correlations in the feedforward input, while the recurrence row arises from the recurrent connections across the populations. **C:** Contributions of paths of given order to networks (left) and the total correlation (right) for the parameters *W*_*EE*_ = 0.25 and *c* = 0.65 (star from panel (A)). All panels: *α* = 0.1.

To better understand the underlying mechanisms responsible for these higher correlations within this parameter regime (i.e., to the right of the pink line in Fig. 2A), we perform a pathway expansion of the covariance matrix Eq. 2. Since the steady state emitted by the system in Eq. 1 is stable, the term (**I**−**W**)^*−*1^ can be expanded as a series. This allows us to write Eq. 2 as (see Methods 5.3)

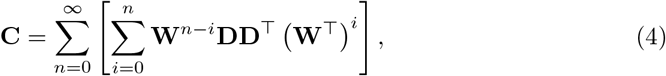

where each term in the inner sum corresponds to an *n*^th^-order path through the network. Writing out the first three terms of this sum for the cross-covariance yields

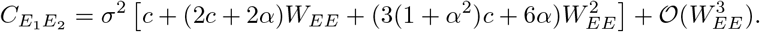

Rewriting this equation as

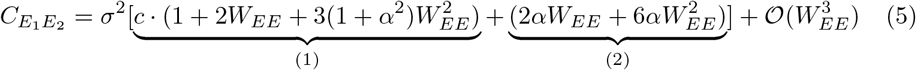

reveals that each term contributing to this cross-covariance can be thought of as arising from one of two sources: 1) inherited from the shared correlated input and dependent on the parameter *c* (Fig. 2B, top), and 2) purely arising from the recurrent connections (Fig. 2B, bottom). We note that the ‘propagation’ of the inherited contribution to higher-order paths does not only rely on the *E*_1_ ↔ *E*_2_ connections (proportional to *α*^*n*^*c*). This is because the correlated activity is fed directly into each subpopulation at the 0^th^ order, from which it can propagate into higher-order paths via self loops contained within each population. We emphasize that eliminating the 0^th^ order term (i.e., setting *c* = 0) eliminates all contributions from the inherited global source.

We now utilize this pathway expansion to compare the contributions from feedforward and recurrent mechanisms to the net cross-covariance for an example point lying in the highly correlated regime (Fig. 2A, star; *W*_*EE*_ = 0.25 and *c* = 0.65). We first note that this series converges quickly and only a few paths significantly contribute to the total correlation (Fig. 2C). The convergence of this series depends directly on the largest eigenvalue of **W** (Methods 5.3), namely

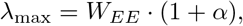

which is small for our choice of parameters. Our numerical results also illustrate that the contribution from the inherited source largely dominates at each order (Fig. 2C; left), and contributes ∼ 90% of the total cross-correlation (Fig. 2C; right). These results hold qualitatively across this parameter regime, and lead us to conclude that it corresponds to a model in which large shared input fluctuations explain the heightened correlations between the separate *E* populations. Taken together, we characterize this solution which exhibits enhanced *E*_1_ − *E*_2_ correlations as a *feedforward inheritance model*.

However, under the condition where the shared input fluctuations are small, we still lack a potential mechanism for significant positive correlations. To surmount this shortcoming, we first need to extend our model to also include inhibitory populations.

### 2.4 Weak recurrent excitation with global inhibition

Parsimoniously, we begin by modelling inhibition as a single global population, consistent with observations that inhibition simply connects densely and non-specifically within cortex (Hofer et al., 2011; Packer and Yuste, 2011) (Fig. 3A). To understand the effect of inhibition in this circuit, we explore how the strength of recurrent inhibitory connections (*W*_*EI*_ *<* 0 and *W*_*IE*_ *>* 0) shape correlations between the excitatory populations in the case when *c* = 0. Assuming *W*_*EE*_ remains weak (i.e., *W*_*EE*_ *<* 1), we find a large portion of the parameter regime yields negative cross-correlations (Fig. 3B; purple region). However, there is a region that satisfies our correlation condition, namely the dark green region that corresponds to strong *I* → *E* and weak *E* → *I* connections.

**Figure 3.**
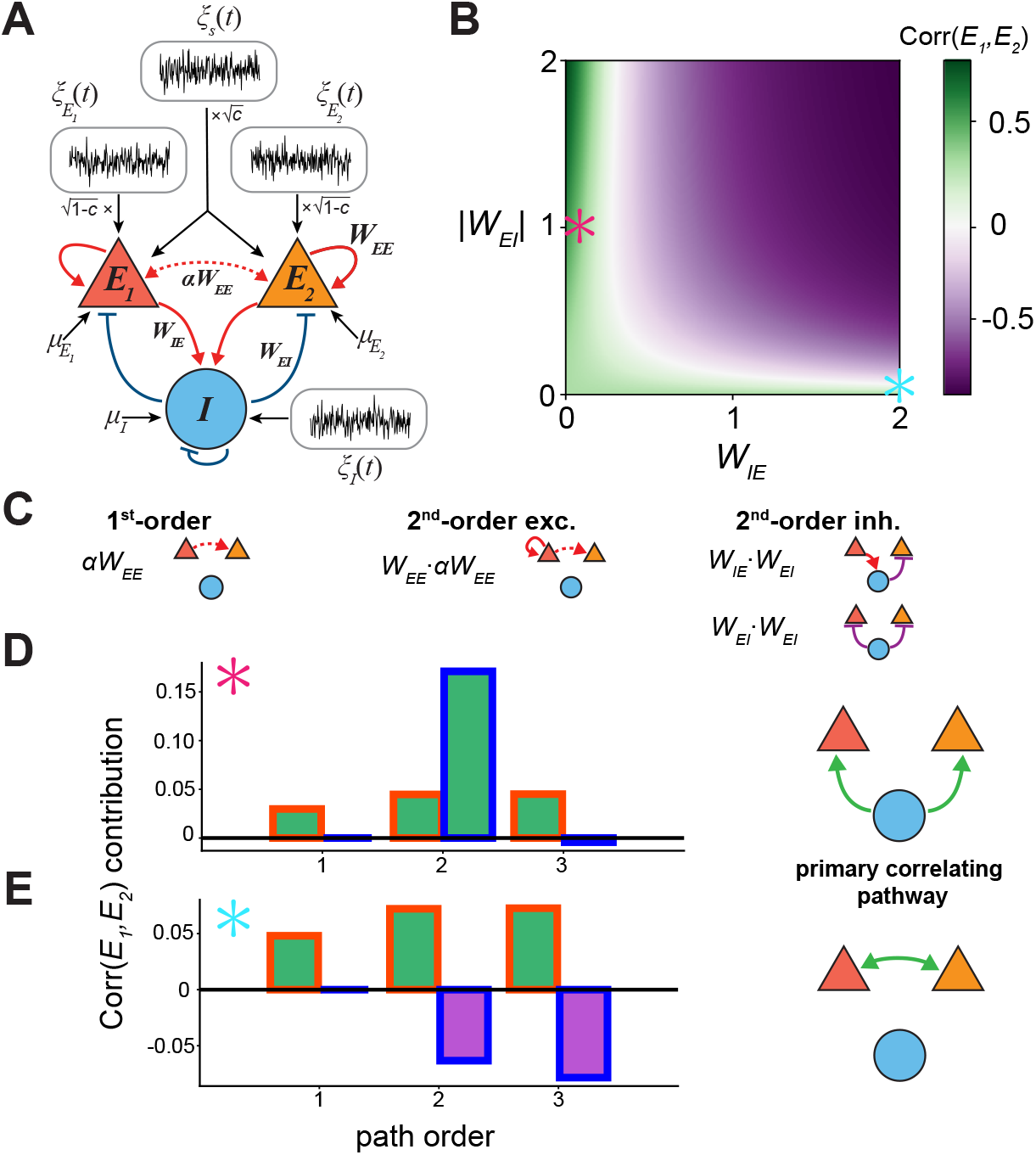
Weakly coupled network. **A:** Network model schematic as in Figure 1C. Blue lines indicate recurrent inhibitory connections. **B:** Corr(*E*_1_, *E*_2_) as a function of |*W*_*EI*_ | and *W*_*IE*_. **C:** Illustrations of first and second order paths. **D, E:** (Left) Contributions of *E* (red outlined bars) and *I* (blue outlined bars) to the net Corr(*E*_1_, *E*_2_). (Right) Schematic of dominant correlating pathway. Colored stars denote locations in B. Red star: *W*_*EI*_ = − 1, *W*_*IE*_ = 0.07; blue star: *W*_*EI*_ = − 0.05, *W*_*IE*_ = 2. For all panels *α* = 0.15.

We again make use of a pathway expansion of Eq. 2 to help decipher this observation, this time accounting for the new inhibitory pathways (Fig. 3C). Writing out the expansion to second order in *W* yields

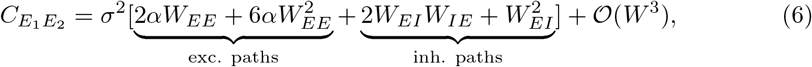

where we have noted the terms involving only the excitatory components and terms which involve paths through the inhibitory population. We first observe that contributions to the cross-covariance due to the excitatory subnetwork at each order are the same as the previous network without the inhibitory connections (Eq. 5 for *c* = 0). This leads us to decompose the total covariance into an excitatory component and an inhibitory component (neglecting the 𝒪 (*W* ^3^) terms in Eq. 6)

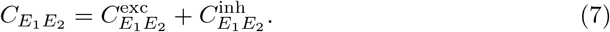

As Eq. 6 suggests, depending on the strength of the underlying inhibitory connections, 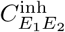 can either be positive (positively correlating the excitatory subpopulations; Fig. 3B, along *W*_*EI*_ axis) or negative (anti-correlating the subpopulations; Fig. 3B, purple region). By contrast, 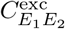 is clearly bounded below by zero.

Specifically, Eq. 6 reveals a ‘tug of war’ that can arise early on in the pathway expansion between the *E* → *I* → *E* (i.e., *W*_*EI*_*W*_*IE*_ *<* 0) and the *I* → *E* (i.e., 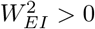) inhibitory pathways. Choosing |*W*_*EI*_ | *> W*_*IE*_ ≈ 0, we find that the positive term dominates, and the the inhibitory population acts as a strong correlator of excitatory activity (Fig. 3D). We term this an *inhibitory inheritance model* by analogy to the feedforward inheritance model described above.

On the other hand, when *W*_*IE*_ *>* |*W*_*EI*_| ≈ 0, the negative term dominates, leading the inhibitory population to weaken the strength of cross-correlations. In this case, the primary correlating source across the excitatory populations are the weak *E*_1_ ↔ *E*_2_ connections (Fig. 3E). But as we noted previously (Fig. 2), this pathway alone is incapable of yielding high cross-correlations without strongly correlated feedforward input.

The regime of weakly coupled neural populations thus permits two solutions for correlating *E*_1_ and *E*_2_ to a sufficiently high degree, both of which can be characterized in terms of inheritance models. Namely, enhanced positive correlations can be inherited from outside sources or from local recurrent inhibition. Nevertheless, the ambiguity in the former solution and the fine-tuning required to achieve the latter solution push us to uncover a more robust mechanism.

### 2.5 Strong recurrent excitation with global inhibition

Up to this point, by virtue of our assumption that the recurrent excitatory coupling is weak the stability of the equilibrium point was independent of the inhibitory currents. Such a network is commonly referred to as a non-inhibition-stabilized network (non-ISN) (Tsodyks et al., 1997; Ozeki et al., 2009; Sadeh and Clopath, 2021) (see Appendix for additional details). However, recent experimental evidence suggests that mouse cortex operates in the ISN regime, where strong recurrent excitation is tracked and balanced by strong inhibitory feedback (Adesnik, 2017; Sanzeni et al., 2020). Since the ISN regime is known to exhibit sometimes perplexing dynamics, such as the well-studied paradoxical effect (Tsodyks et al., 1997), it is initially unclear how shifting into this parameter regime will shape the correlations under investigation.

In view of this, we now strengthen the recurrent excitatory connections *W*_*EE*_ such that our model network lies in the ISN regime. Performing a similar analysis as before (i.e., fixing *W*_*EE*_ and *W*_*II*_, while varying *W*_*EI*_ and *W*_*IE*_) and assuming that the feedforward inputs are uncorrelated (*c* = 0), we find results that at first glance appear familiar (Fig. 4A). Namely, a portion of the parameter regime results in negative correlations (purple region), with a narrow parameter regime yielding positive correlations (green region). However, unlike the previous network, these correlations are much larger across this band of parameter values, approaching unity as the system loses stability due to the inhibitory feedback becoming too weak to be able to balance out the strong excitation (gray and red-hatched region).

**Figure 4.**
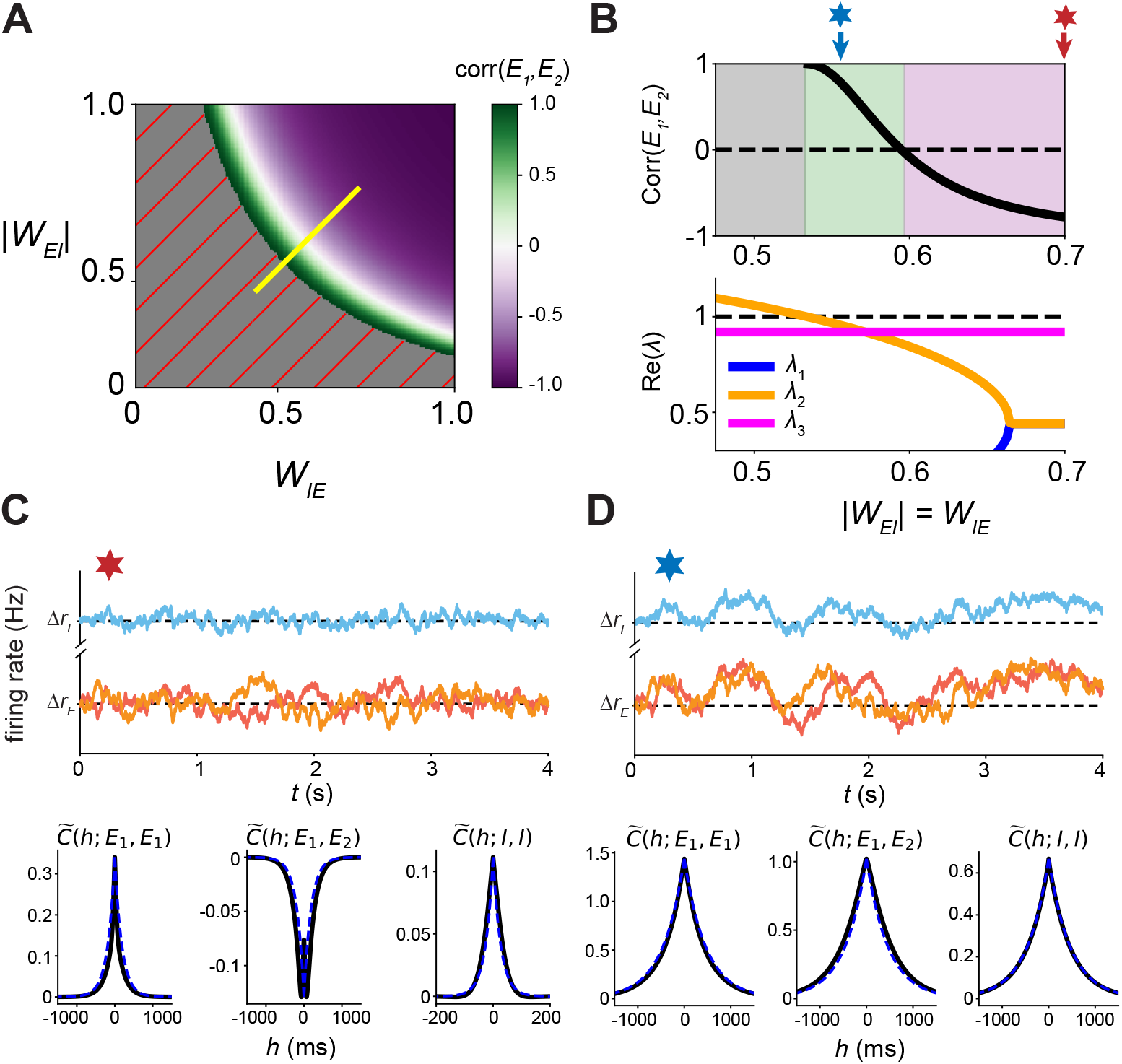
Global inhibition in ISN regime. **A:** Corr(*E*_1_, *E*_2_) as a function of *W*_*EI*_, *W*_*IE*_ with *c* = 0. **B:** Top: Corr(*E*_1_, *E*_2_) along the yellow path in A. Gray region: unstable; green region: positive correlations; purple region: negative correlations. Bottom: eigenvalues of the circuit along the yellow path in A. **C, D:** Top: example rate traces (colors as in Fig. 3B). Bottom: auto- and cross-correlation functions computed numerically (black) and theoretically for the dominant timescale (blue dashed). Stars indicate parameter values shown in B. Here, *α* = 0.2.

Unlike the non-ISN regime, where the weak recurrent excitatory connections corresponded with small eigenvalues and quick convergence in our path-expansion, here the eigenvalues of the system lie much closer to the boundary separating stability from instability. As result, many more terms are needed before the series in Eq. 4 converges, complicating its interpretation. Instead, we seek to understand the mechanism driving these high correlations by exploring their apparent connection to the system’s stability.

We start by considering the slice of the parameter space where |*W*_*EI*_| = *W*_*IE*_ that captures the system’s transitions from negative correlations to positive correlation to instability (Fig. 4A, yellow line; Fig. 4B, top). Analysis of the eigenvalues of **W** reveals a pair of eigenvalues (*λ*_1_ and *λ*_2_) dependent on the strength of inhibitory connections and another eigenvalue that remains constant (and close to one) along this parameter slice (*λ*_3_ = *W*_*EE*_(1 − *α*)) (Fig. 4B, bottom). Interestingly, we find that decay for the stationary autocovariance function for the inhibitory population (Fig. 4D and 4E bottom; see Eq. 9 in Section 5) is well approximated by

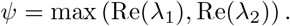

From this link, we see that when |*W*_*EI*_| = *W*_*IE*_ is large, then *ψ* is small, meaning the timescale of inhibition is fast. This allows the inhibitory population to rapidly and effectively cancel the net excitatory inputs (Fig. 4C, top). We observe that in this parameter regime, Δ*r*_*I*_ remains small, while 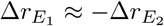, leading to strong negative correlations between *E*_1_ and *E*_2_. As |*W*_*EI*_| = *W*_*IE*_ decreases, *ψ* increases towards one, which slows down the inhibitory timescale (Fig. 4D, top). This slower cancellation of the excitatory currents allows for larger deviations away from baseline for all neuronal populations. However, since the system is still stable, we observe that the populations co-vary together, leading to correlated excursions in the rates.

In total, the ISN regime yielded a more robust set of parameter values corresponding to high correlations across the segregated excitatory populations than the non-ISN regime observed previously. However, even in this improved scenario, the viable parameter regime is still limited to a relatively thin band, and further, this band lies precariously close to regions of instability.

### 2.6 Strong recurrent excitation with clustered inhibition

The fine tuning required to capture large Corr(*E*_1_, *E*_2_) despite having weak *E*_1_ ↔ *E*_2_ coupling (*α <<* 1) for both the purely excitatory and global inhibitory networks places doubt on these mechanisms being operative in real neuronal circuits. In this section we hypothesize that if the sources of inhibition for each excitatory subpopulation are similarly clustered, then this decoupling of inhibition may permit a larger stable region of positive correlations in the ISN regime, largely by limiting the effects of the anticorrelating *E*_1_ → *I* → *E*_2_ and *E*_2_ → *I* → *E*_1_ pathways.

We implemented inhibition to be co-clustered with the excitatory subpopulations by separating the inhibitory population in two, with *I*_1_ and *I*_2_ corresponding to the respective excitatory populations *E*_1_ and *E*_2_ (Fig. 5A). In this case, each *E*_*i*_*/I*_*i*_ cluster constitutes an ISN (*i* = 1, 2). The model contains no inter-population connections except those between *E*_1_ and *E*_2_, and without any source of shared input correlations (*c* = 0). We have again assumed symmetry in the connection strengths such that the pairs (*E*_1_, *I*_1_) and (*E*_2_, *I*_2_) are identical in their connectivity and dynamics.

**Figure 5.**
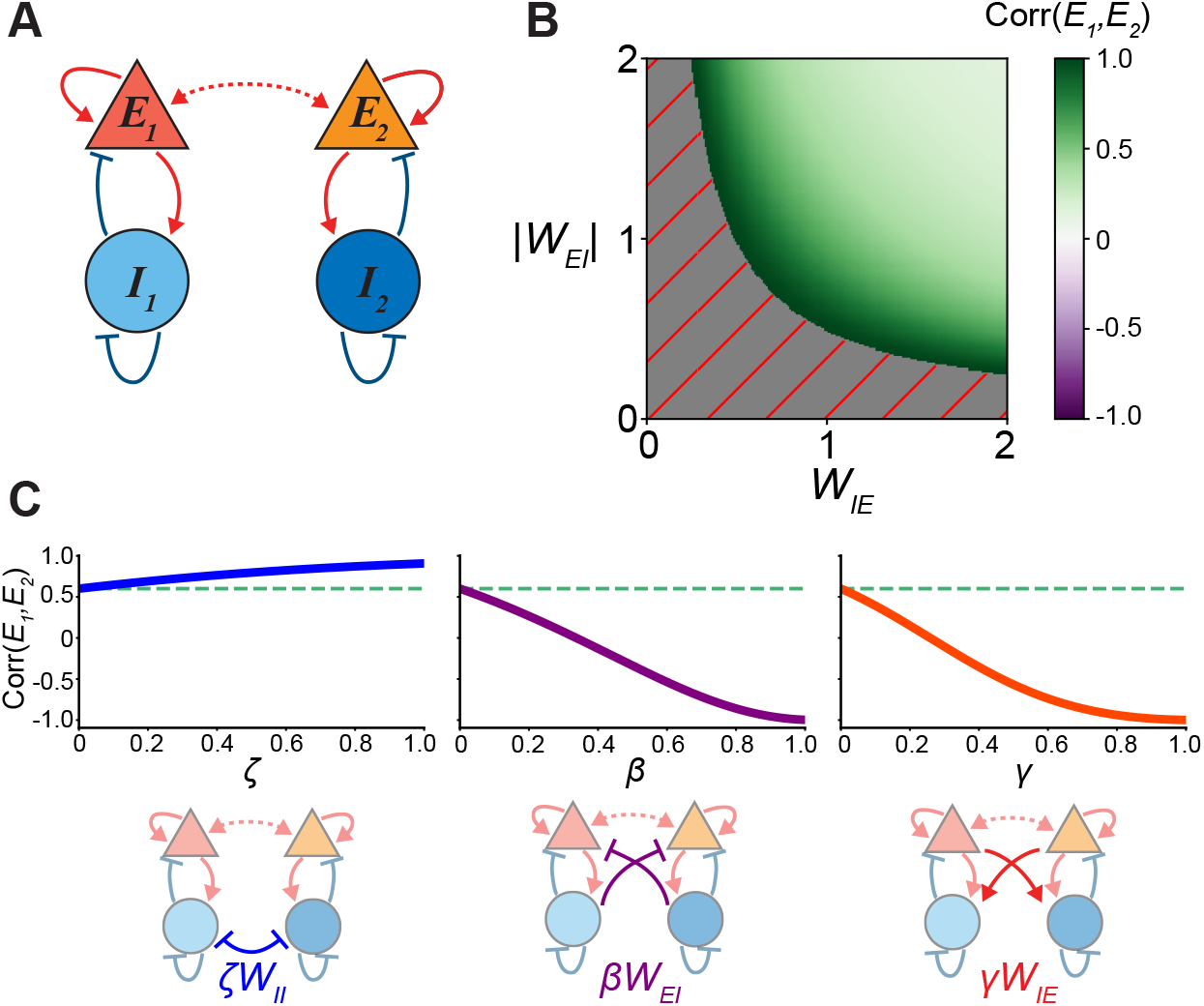
Segregated *I* subpopulations produce robust positive correlations. **A:** Model schematic. Input structure is consistent with Fig. 3A but omitted for clarity. **B:** Corr(*E*_1_, *E*_2_) as a function of *W*_*EI*_, *W*_*IE*_ with *c* = 0. **C:** Corr(*E*_1_, *E*_2_) as a function of added connections between *I*_1_, *I*_2_ (left); *I*_1_ → *E*_2_ and *I*_2_ → *E*_1_ (middle); *E*_1_ → *I*_2_ and *E*_2_ → *I*_1_ (right). Added connections *W*_*ij*_ are initialized to the same as elsewhere in the network, and scaled by: *ζ, I* ↔ *I*; *β, I* → *E*; *γ, E* → *I*. Dashed turquoise line denotes *ζ, β, γ* = 0, *W*_*EI*_ = *W*_*IE*_ = 1.

If we again fix *W*_*EE*_, *W*_*II*_ and proceed by exploring the space of *W*_*EI*_, *W*_*IE*_ connections, we find that this network structure now yields a robust region in which correlations are strong and positive (Fig. 5B, green). This can only be due to the strong dynamic recruitment of the inter-population connection *αW*_*EE*_. This result emphasizes three important points. First, that there exists a large space of connection parameters in which our criteria (large Corr(*E*_1_, *E*_2_) and *α <<* 1) may be met. Given the heterogeneity of neural circuits and plasticity of connections within cortex this parametric result is much more satisfying than a fine-tuned solution like that required in the model with global inhibition (Sections 2.4, 2.5). Second, this result does not depend on the presence of external correlated fluctuations. Thirdly, this result is robust to the presence of external correlated input noise as it would only further amplify the observed correlations.

A natural question is whether incorporating inter-population *E*_*i*_ ↔ *I*_*j*_ or *I*_*i*_ ↔ *I*_*j*_ connections would affect this result. We therefore considered fixed values of *W*_*EI*_, *W*_*IE*_, and *W*_*II*_, and introduced scaling parameters *β, γ, ζ*, respectively, to adjust the between-population strengths of each connection (Fig. 5C). We found that only *W*_*II*_ *>* 0 was able to further enhance correlations above the value we found when *ζ* = 0 (Fig. 5C, left). In contrast, any non-zero values of *β, γ* only reduced correlations (Fig. 5C, middle and right). This is due to the same mechanism discussed previously in which strong excitatory recruitment of inhibition induces anti-correlations between the populations. Further, this same relationship also held when *W*_*EI*_, *W*_*IE*_, *W*_*II*_ were co-varied. Only *I* → *I* connections served as a correlating force; all others induced a reduction in correlations (AFig. 2). Hence, we conclude that while inhibition can be promiscuously connected with other inhibitory units, it must be strongly co-clustered with excitatory subpopulations and sparse in its connectivity with other excitatory subpopulations to yield the significant positive inter-population excitatory correlations observed in Kim et al. (2018b).

## 3 Discussion

In this study we sought to uncover possible neural circuit mechanisms underpinning the experimental observation that pyramidal neurons projecting to different downstream targets connect with a much lower probability than random pairs of excitatory neurons, yet still exhibit correlated variability that is almost as large as the rest of mouse V1 (Kim et al., 2018b). Notably, the magnitude of these correlations is much stronger than would be predicted given their weak connectivity. We found that a model with global inhibition resulted in highly constrained regions in which the data could be matched, encompassing two distinct solutions. In the case of weak network coupling, positive correlations resulted from two forms of *inheritance model* : either *I* → *E* connections induced increased correlated activity through *I* affecting both excitatory populations in the same way, or an unobserved external source of strong correlations fed these fluctuations across both *E* units. When connectivity strengths grew, placing the circuit in an inhibition-stabilized (ISN) regime, the network needed to live right at the edge of stability to observe positive correlations. By contrast, we found that a more generally robust solution in the ISN regime could be achieved by splitting the inhibitory population into two separate subpopulations co-clustered with one of the excitatory subnetworks.

We argue that, on the basis of this robustness, our results thus predict that inhibition should cluster together with excitation in mouse sensory cortex with a specificity that mirrors that of the excitatory connectivity. The other inferred models by contrast depend upon narrow parameter regimes to capture experimental observations. This fragility would require significant constraints on the properties of neural circuits. Yet, connections are plastic, connection strengths are heterogeneous, and neuron properties are affected by neuromodulation (Turrigiano, 2008; Marder, 2012). Given this stochasticity in the circuit structure itself, a fine-tuned solution is unlikely to capture the data.

Rigorous experimental validation of our model predictions could be obtained through physiological or connectomics experiments which specifically target the relationship between excitatory projection neurons and local inhibitory neurons. While it is well-appreciated that inhibitory interneurons are very diverse in physiology and connectivity (Tremblay et al., 2016), we did not explicitly model this diversity in our study. Nevertheless, we anticipate that parvalbumin (PV)-positive cells may display the identified signatures of our *I* units, as they appear to play a critical role in stabilizing excitatory activity (Bos et al., 2020). Recent experimental evidence appears to support this claim from the perspective of stimulus tuning: while PV cells connect with most nearby pyramidal neurons, they were found to more strongly connect with those whose tuning properties they share (Znamenskiy et al., 2018).

Recent theoretical work has argued that *E*/PV assembly formation requires plasticity from both *E* → PV and PV → *E* connections (Mackwood et al., 2021). This bidirectionality could result in local, winner-take-all effects in *E* ↔ *I* connectivity as any discrepancies in functional response properties between nearby pyramidal cells will bias the PV connectivity. This could result in the more specific co-clustering of inhibition we predict. Motivated by these results, a potential indirect way to differentiate between the global and clustered inhibition models would be to record activity of AL- and PM-projecting neurons together with inhibitory interneurons.

Comparison of their respective tuning functions could suggest whether the inhibitory cell is biased in its connectivity (by extension of Znamenskiy et al. (2018)). Indeed, Najafi et al. (2020) recently argued for co-clustered excitation-inhibition in the context of posterior parietal cortex decision circuitry on the basis of neural response properties. Furthermore, in mouse visual cortex, it has been shown that PM and AL exhibit distinct functional representations with some overlap (Andermann et al., 2011), consistent with the tuning properties of V1 projection neurons (Kim et al., 2018b). Of course, it is possible that inhibitory-excitatory interactions may span a continuum between the global and clustered motifs identified here. This raises the possibility that heterogeneity in inhibitory connectivity motifs at small spatial scales may explain heterogeneity in pairwise covariance between AL- and PM-projecting pyramidal cells.

A central issue in the extension of our results concerns the dynamical regime of cortex, a topic which has received a significant amount of attention lately (Ahmadian and Miller, 2021; Morales et al., 2021; Huang, 2021). One question concerns whether intracortical interactions are strong enough to require inhibition as a key stabilizer of activity, that is, whether sensory cortex is an inhibition-stabilized network (Tsodyks et al., 1997; Sadeh and Clopath, 2021). Theoretical work predicts that in this regime the ratio of excitatory to inhibitory input drive to a neuron decreases with increasing stimulus intensity (Rubin et al., 2015). Recent experimental evidence from recordings of mouse primary visual cortex supports this claim (Adesnik, 2017). Another study used optogenetic perturbation of inhibitory neurons across mouse cortex to test for inhibition-stabilization without sensory stimulation, finding evidence that all considered cortical regions operate as an ISN (Sanzeni et al., 2020).

Given this evidence for an ISN regime, a second question regards whether the network dynamics are poised near a change in stability. In our model, loss of stability would result in large positive correlations through a slowing down of the dynamics (Fig. 4). Analysis of large-scale recordings in mice has suggested that cortex may in fact live close to an instability (Morales et al., 2021). This could suggest that either the global or clustered inhibition model in an ISN regime may explain the data. Together with the foregoing evidence that PV and *E* neurons sharing tuning properties connect more strongly, we argue that this further supports a model of co-clustered inhibition.

Other mechanisms by which correlations can grow near a change in stability have been identified in previous studies. Ginzburg and Sompolinsky (1994) observed that near a bifurcation - in their case, a saddle node or Hopf - correlations in a weakly connected network grow from 𝒪 (1*/N*) to near 𝒪 (1) where *N* is the network size, together with a slowing down in the dynamics. Darshan et al. (2018) derived conditions on what they term the interaction matrix (similar to our **W** matrix) under which correlations are amplified without critical slowing down. These network models thus suggest distinct mechanisms by which our results could be extended to spatially-distributed spiking network models. Additionally, Litwin-Kumar and Doiron (2012) studied the effect of clustered connectivity in balanced spiking networks on the structure of correlations, however this work did not compare across-cluster to within-cluster correlations. Rosenbaum et al. (2017) did consider a structure similar to our three-population global inhibition motif, demonstrating that, consistent with our conclusions, a spatially distributed spiking neural network with distinct subpopulations would show close to zero correlations on average due to strong positive correlations within a cluster and large negative correlations between the two clusters. Yet it remains for future work to determine the precise parametric values to recapitulate our results in spiking neural network models.

Our work can be seen as a case study of a particular network structure in the context of the theoretical investigation of dynamics on graphs (that is, a collection of nodes and edges). In general, graphical analysis has been used in a wide range of neuroscientific applications, from the determination of fixed points of dynamics (Morrison and Curto, 2019) to network controllability (Kim et al., 2018a). In relating connectivity motifs (elements of **W** and their combinations) to correlation structure in the circuit, our approach relates to a more general mathematical concept of relating process motifs on networks to underlying structure motifs of the graph (Schwarze and Porter, 2021).

Ultimately, this work demonstrates how ostensibly straight-forward observations of connectivity and response properties from cortical cells have the capacity to lend fruitful insight into the structural and dynamical regimes of cortex, which are critical to further understanding of information processing in the brain.

## 4 Acknowledgments

A.N., M.P.G., and G.H. thank the Simons Foundation SCGB Undergraduate Research Fellowship (SURF) for fostering this collaboration. A.N. was funded by Simons Foundation SURF. G.H. was supported by the Burroughs Wellcome Fund’s Career Award at the Scientific Interface. B.D. is supported by National Institutes of Health (NIH) (grant no. 1U19NS107613-01, R01EB026953), Vannevar Bush faculty fellowship (no. N00014-18-1-2002) and the Simons Foundation Collaboration on the Global Brain.

## 5 Methods

### 5.1 Firing rate model

As done previously (Kanashiro et al., 2017; Getz et al., 2022), we consider the firing rate dynamics of neuronal populations *A* given by the following

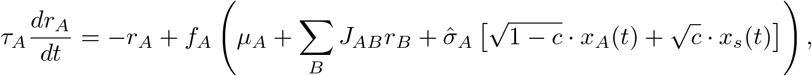

where *τ*_*A*_ is the time constant, *μ*_*A*_ is a constant stimulus drive, and *J*_*AB*_ is the strength of connections from population *B* to *A*. The stochastic processes *x*_*A*_(*t*) and *x*_*s*_(*t*) represent private and shared global fluctuations, respectively. Each is taken to be the limiting process from

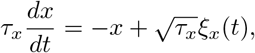

for *τ*_*x*_ → 0, with ⟨*ξ*_*i*_(*t*)⟩ = 0 and ⟨*ξ*_*i*_(*t*)*ξ*_*i*_(*t*^*′*^)⟩ = *δ*(*t* − *t*^*′*^). Intuitively, one may think of *x*(*t*) as a “smoothed” white noise process (Kanashiro et al., 2017). The parameter *c* ∈ [0, 1] scales the proportion of shared noise relative to private noise, while 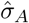 represents the total intensity of the fluctuations.

We assume that the system of equations has an equilibrium point at *r*_ss_, and that the noise is weak enough so that the fluctuations about this equilibrium (Δ*r* := *r* − *r*_ss_) can be approximated by

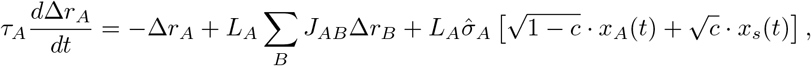

where 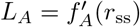 is the gain of population A at the equilibrium point. We define the effective coupling as *W*_*AB*_ := *L*_*A*_*J*_*AB*_ and 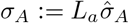, and approximate *x*_*A*_(*t*) and *x*_*s*_(*t*) as independent, zero-mean Gaussian processes *ξ*_*A*_(*t*) and *ξ*_*s*_(*t*) satisfying ⟨*ξ*(*t*)*ξ*(*t*^*′*^)⟩ = *δ*(*t* − *t*^*′*^). This yields Eq. 1, which in matrix form can be written as

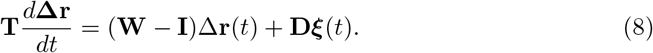

For notational simplicity, throughout we will assume unit time constants *τ*_*A*_ = 1, so that **T** = **I**. For example, in the case of two excitatory populations and one inhibitory population {*E*_1_, *E*_2_, *I*} the matrices are

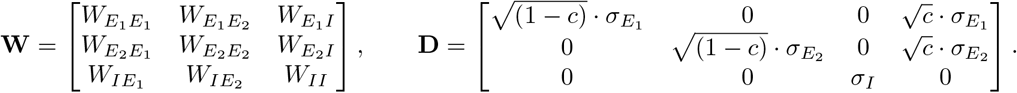

The network structure is determined through the weight matrix **W**. Since we are explicitly interested in segregated excitatory populations, we consider weak the cross-population connections and set

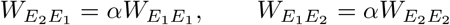

for *α* ∈ (0, 1). The two excitatory populations, *E*_1_ and *E*_2_, are increasingly disconnected as *α* → 0. To obtain analytical expressions and constrain the searchable parameter space, we assume various symmetries in the network connectivity. Specifically, we consider the following forms for connectivity matrices for the two (Fig. 1A), three (Fig. 3A) and four (Fig. 4A) population models:

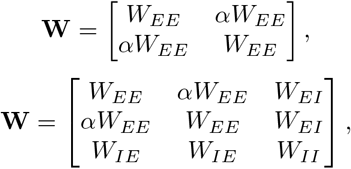

and

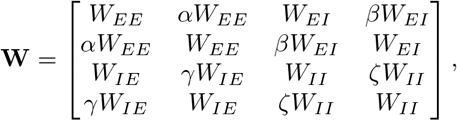

where *β, γ, ζ* ∈ (0, 1).

### 5.2 Covariance calculation

The autocovariance function for the OU process defined in Eq. 8 is given by

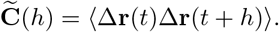

Let **M** = **W** − **I** and define 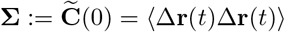 as the stationary covariance matrix. Then **Σ** is obtained as the solution to the *Lyapunov equation* −**MΣ** + Σ(−**M**)^⊤^ = **DD**^⊤^ (Gardiner, 2009). *It follows that*

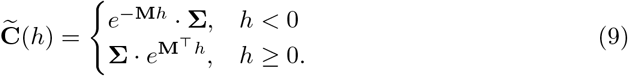

Integrating 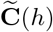 in each element over long times *h* yields the following compressed form for the *long-time covariance matrix* **C**

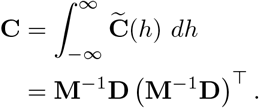

If 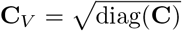, then the correlation matrix is obtained

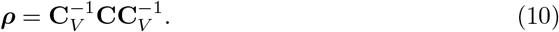

### 5.3 Path expansion

If the spectral radius *s*(**W**) = max{|*λ*_*i*_| : *λ*_*i*_ is an eigenvalue of **W**} *<* 1, then **M**^*−*1^ has a convergent series representation

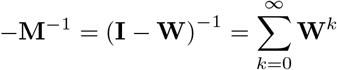

known as a *Neumann series* (Einsiedler et al., 2017). Intuitively, one may think of the Neumann series as a matrix analogue of the familiar geometric series. Under this representation, the long-time covariance matrix is

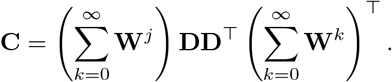

It is useful to rewrite this expansion as

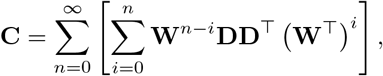

where the terms in the inner sum can be interpreted as contributions due to *n*^th^-order paths through the network (Trousdale et al., 2012; Pernice et al., 2011).

If the outer sum converges quickly, the covariance matrix can be approximated as

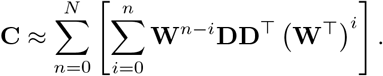

The rate of convergence of this approximation depends on the magnitude of *s*(**W**). In particular, the closer *s*(**W**) is to 0, the faster the terms shrink. Consider the *N* -th order terms of this approximation,

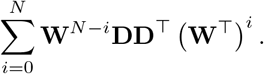

If || · || is the operator norm, then

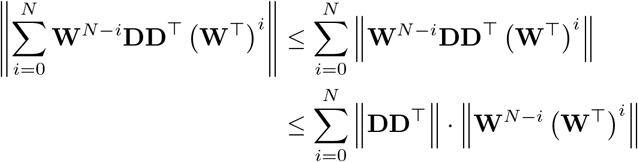

Diagonalize **W** and write **W** = **PΛP**^*−*1^, where **Λ** is a diagonal matrix of eigenvalues of **W**. It follows that

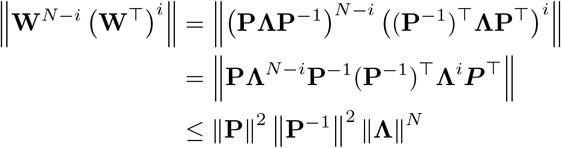

Then

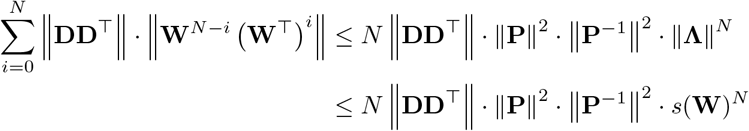

This bound shrinks quickly as *N* → ∞ if *s*(**W**) is small (≪1), as is the case when the system is in the weakly coupled regime.

#### 5.3.1 Path expansion for weakly coupled *E*_1_ ↔ *E*_2_

In Fig. 2B we illustrate this quick convergence by showing the first three terms of this sum, namely

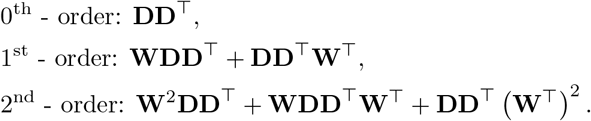

Using these terms, the cross population covariance can be approximated by Eqn. 5 in the main text.

We note that for a *n*^th^-order path, we multiply on the left and right by 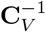 to obtain path contributions to the correlation matrix. In particular, we are interested in the contributions to 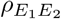 (that is, the element ***ρ***_1,2_ of Eqn. 10).

### 5.4 Parameters & Simulations

All relevant code will be made available at the author’s github upon publication. Simulations were performed using an Euler-Maruyama scheme with time constants *τ*_*E*_ = *τ*_*I*_ = 15 msec, *dt* = 0.01 msec.

**Table 1.**
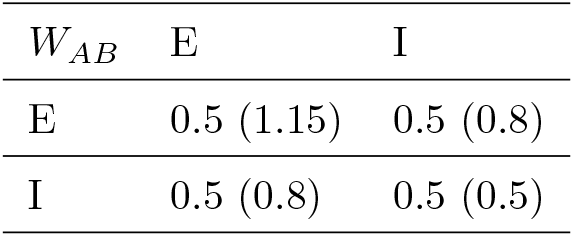
Strength of connections from pop. *B* **(columns) to** *A* **(rows) for the weakly (strongly) coupled model**.

**Table 2.**
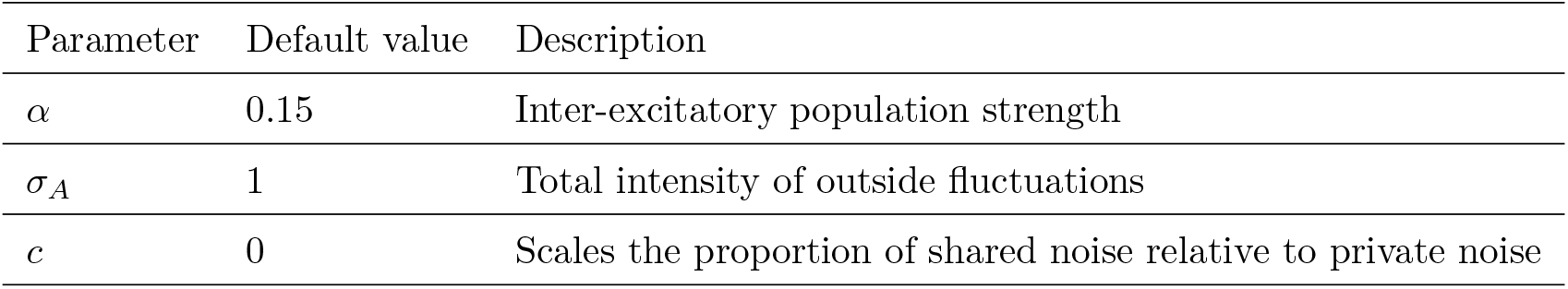
Default parameter values. Changes to any parameter are indicated in the figure caption.

## 6 Supplemental Information

### 6.1 Network stability

The deterministic version of Eqn. 8 (Fig. 1A)

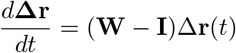

is *asymptotically stable* if the eigenvalues *λ*_*i*_ of **W** − **I** satisfy ℜ [*λ*_*i*_] *<* 0, meaning that a perturbation of the excitatory rates is quenched and rates are returned to their steady-state values (AFig. 1B, top) (Wiggins, 2003). An equivalent condition for stability is if the eigenvalues *λ*_*i*_ of **W** satisfy ℜ {*λ*_*i*_} *<* 1. We say that a network is stable if it admits a stable equilibrium solution, otherwise we say the network is unstable.

**Supplemental Figure 1.**
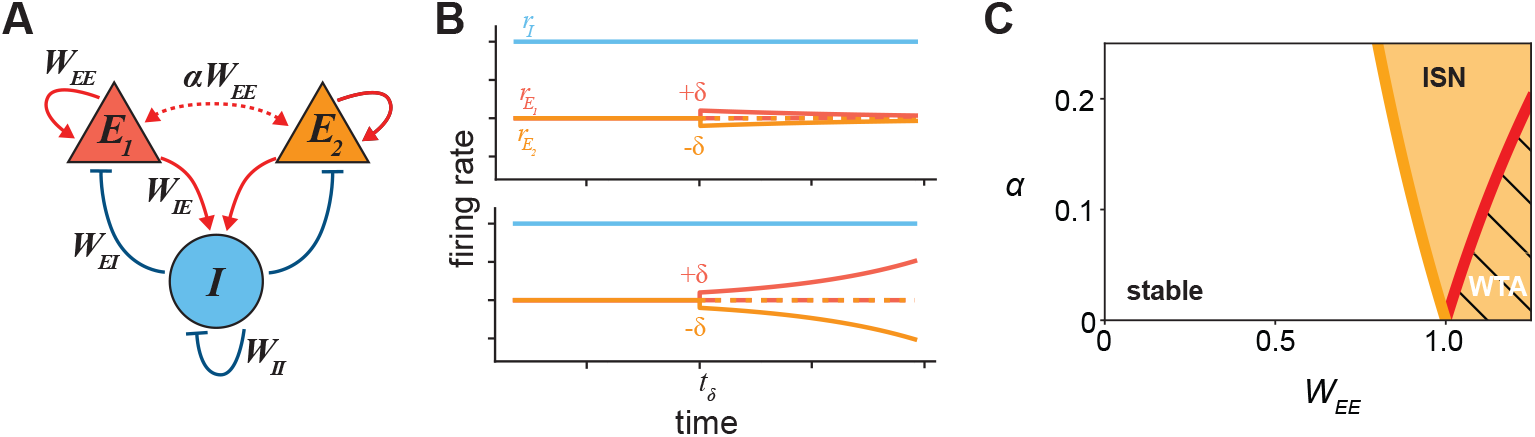
Dynamical regimes and limitations on *α*. **A:** Network schematic. **B:** Illustrations of a small change in the input +*δ* to *E*_1_ and −*δ* to *E*_2_. Top: stable network regime; bottom: unstable (winner-take-all) regime. **C:** *W*_*EE*_ − *α* space. Yellow region: inhibition-stabilized (ISN); black hatched region: winner-take-all (unstable). Solid yellow line: *W*_*EE*_ = 1*/*(1 + *α*). Solid red line: *α* = 1 − 1*/W*_*EE*_. Parameters as in Fig. 5. *α* changed to 0.1 for the unstable regime in B.

#### 6.1.1 The inhibition-stabilized network (ISN)

A linear network is an *inhibition stabilized network* (ISN) (Ozeki et al., 2009) if it satisfies two conditions:

1. The network is unstable in the absence of (dynamic) feedback inhibition,
2. The network is stable with sufficiently strong inhibition.

We consider the conditions under which the global inhibition motif (i.e., two excitatory populations with one shared inhibitory population) is an ISN. The corresponding weight matrix is

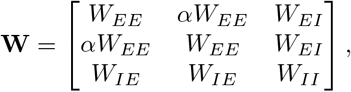

which has eigenvalues

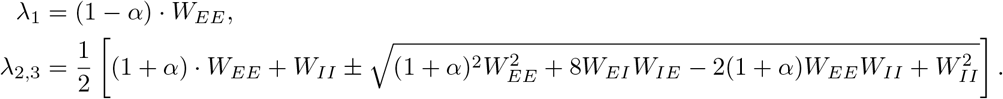

We note that *λ*_1_ does not depend on any of the inhibitory connections. As a result, if *λ*_1_ = (1 − *α*) · *W*_*EE*_ *>* 1 the system is unstable and inhibition is unable to stabilize it, so we necessarily require (1 − *α*) · *W*_*EE*_ *<* 1. On the other hand, *λ*_2,3_ do depend on the inhibitory connections. Absent feedback inhibition (i.e., *W*_*EI*_ = 0) these eigenvalues become

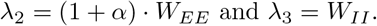

In this work the latter is always less than 1. Meanwhile, it is possible to increase recurrent excitation such that *λ*_2_ = (1 + *α*) · *W*_*EE*_ *>* 1. Unlike the previous condition derived with *λ*_1_, we can choose inhibitory parameters *W*_*EI*_, *W*_*II*_ such that this eigenvalue decreases below 1, restoring the stability of the system. Thus, this system lies in the ISN regime when

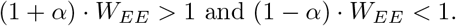

If the first condition is satisfied, while the second condition is violated, the system exhibits winner-take-all dynamics, where one excitatory population increases away from steady state while the second decreases away from it (AFig. 1B, bottom). All three regions (non-ISN, ISN, and winner-take-all) are shown in AFig. 1C. The same constraint to lie in the ISN regime can also be derived for the specific inhibition motif.

### 6.2 Covarying cross-population connections

It is possible that in the segregated *E*_*i*_*/I*_*i*_ subpopulation model (Fig. 5A), covarying cross-population connections might induce synergetic effects different from those observed by adding singular bidirectional connections (Fig. 5C). We tested this numerically by adding pairwise combinations of *E* → *I*, → *I E*, and *I* → *I* (AFig. 2A-C). Only when *ζ* (scaling of *I* → *I*) dominated either *β* or *γ* was an increase in correlations observed; *E* → *I* and *I* → *E* always reduced correlations, consistent with the results presented above.

**Supplemental Figure 2.**
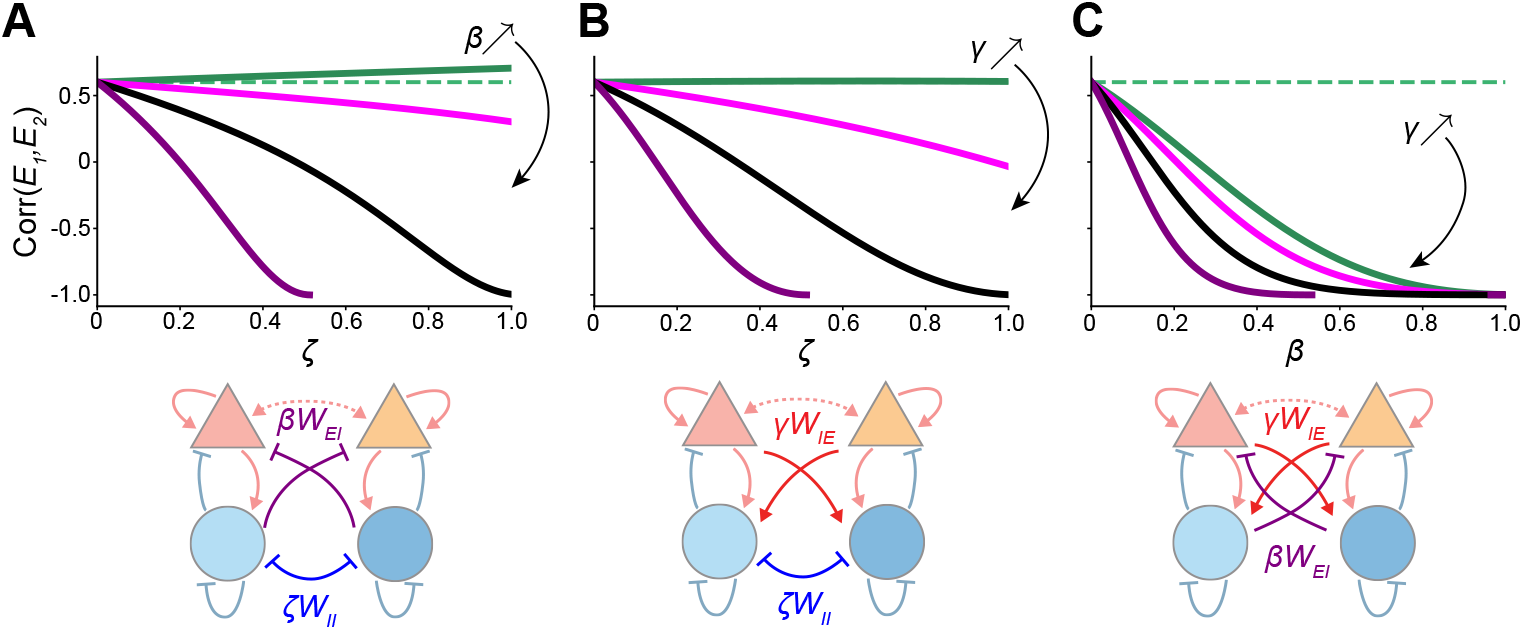
Covarying cross-population connections in segregated ISN. **A:** Top: *Corr*(*E*_1_, *E*_2_) as a function of *ζ*; colored lines indicate different values of *β*. Black lines indicate when *ζ* = *β*. Bottom: network schematics indicating connection weights co-varied in the above plot. Green dashed line indicates value of *Corr*(*E*_1_, *E*_2_) for *W*_*EI*_ = *W*_*IE*_. Parameters as in Fig. 5. **B:** same as (A) for *ζ* and *γ*. **C:** same as (A) for *β* and *γ*.

## References

H. Adesnik. Synaptic mechanisms of feature coding in the visual cortex of awake mice. Neuron, 95(5):1147–1159, 2017.

Y. Ahmadian and K. D. Miller. What is the dynamical regime of cerebral cortex? Neuron, 109(21):3373–3391, 2021.

M. L. Andermann, A. M. Kerlin, D. K. Roumis, L. L. Glickfeld, and R. C. Reid. Functional specialization of mouse higher visual cortical areas. Neuron, 72(6): 1025–1039, 2011.

T. Biswas and J. E. Fitzgerald. Geometric framework to predict structure from function in neural networks. Physical Review Research, 4(2):023255, 2022.

H. Bos, A.-M. Oswald, and B. Doiron. Untangling stability and gain modulation in cortical circuits with multiple interneuron classes. bioRxiv, pages 2020–06, 2020.

M. R. Cohen and A. Kohn. Measuring and interpreting neuronal correlations. Nature neuroscience, 14(7):811–819, 2011.

L. Cossell, M. F. Iacaruso, D. R. Muir, R. Houlton, E. N. Sader, H. Ko, S. B. Hofer, and T. D. Mrsic-Flogel. Functional organization of excitatory synaptic strength in primary visual cortex. Nature, 518(7539):399–403, 2015.

R. Darshan, C. Van Vreeswijk, and D. Hansel. Strength of correlations in strongly recurrent neuronal networks. Physical Review X, 8(3):031072, 2018.

A. Das and I. R. Fiete. Systematic errors in connectivity inferred from activity in strongly recurrent networks. Nature Neuroscience, 23(10):1286–1296, 2020.

B. Doiron, A. Litwin-Kumar, R. Rosenbaum, G. K. Ocker, and K. Josić. The mechanics of state-dependent neural correlations. Nature neuroscience, 19(3):383–393, 2016.

M. Einsiedler, T. Ward, et al. Functional analysis, spectral theory, and applications, volume 104. Springer, 2017.

A. A. Faisal, L. P. Selen, and D. M. Wolpert. Noise in the nervous system. Nature reviews neuroscience, 9(4):292–303, 2008.

C. Gardiner. Stochastic methods, volume 4. Springer Berlin, 2009.

M. P. Getz, C. Huang, and B. Doiron. Subpopulation codes permit information modulation across cortical states. bioRxiv, pages 2022–09, 2022.

I. Ginzburg and H. Sompolinsky. Theory of correlations in stochastic neural networks. Physical review E, 50(4):3171, 1994.

K. M. Hagihara, A. W. Ishikawa, Y. Yoshimura, Y. Tagawa, and K. Ohki. Long-range interhemispheric projection neurons show biased response properties and fine-scale local subnetworks in mouse visual cortex. Cerebral Cortex, 31(2):1307–1315, 2021.

S. B. Hofer, H. Ko, B. Pichler, J. Vogelstein, H. Ros, H. Zeng, E. Lein, N. A. Lesica, and T. D. Mrsic-Flogel. Differential connectivity and response dynamics of excitatory and inhibitory neurons in visual cortex. Nature neuroscience, 14(8):1045–1052, 2011.

C. Huang. Modulation of the dynamical state in cortical network models. Current opinion in neurobiology, 70:43–50, 2021.

T. Kanashiro, G. K. Ocker, M. R. Cohen, and B. Doiron. Attentional modulation of neuronal variability in circuit models of cortex. Elife, 6:e23978, 2017.

J. Z. Kim, J. M. Soffer, A. E. Kahn, J. M. Vettel, F. Pasqualetti, and D. S. Bassett. Role of graph architecture in controlling dynamical networks with applications to neural systems. Nature physics, 14(1):91–98, 2018a.

M.-H. Kim, P. Znamenskiy, M. F. Iacaruso, and T. D. Mrsic-Flogel. Segregated subnetworks of intracortical projection neurons in primary visual cortex. Neuron, 100 (6):1313–1321, 2018b.

H. Ko, S. B. Hofer, B. Pichler, K. A. Buchanan, P. J. Sjöström, and T. D. Mrsic-Flogel. Functional specificity of local synaptic connections in neocortical networks. Nature, 473(7345):87–91, 2011.

A. Litwin-Kumar and B. Doiron. Slow dynamics and high variability in balanced cortical networks with clustered connections. Nature neuroscience, 15(11):1498–1505, 2012.

O. Mackwood, L. B. Naumann, and H. Sprekeler. Learning excitatory-inhibitory neuronal assemblies in recurrent networks. Elife, 10:e59715, 2021.

E. Marder. Neuromodulation of neuronal circuits: back to the future. Neuron, 76(1): 1–11, 2012.

Y. Mishchencko, J. T. Vogelstein, and L. Paninski. A bayesian approach for inferring neuronal connectivity from calcium fluorescent imaging data. The Annals of Applied Statistics, pages 1229–1261, 2011.

G. B. Morales, S. Di Santo, and M. A. Muñoz. Quasi-universal scaling in mouse-brain neuronal activity stems from edge-of-instability critical dynamics. arXiv preprint arXiv:2111.12067, 2021.

K. Morrison and C. Curto. Predicting neural network dynamics via graphical analysis. In Algebraic and Combinatorial Computational Biology, pages 241–277. Elsevier, 2019.

F. Najafi, G. F. Elsayed, R. Cao, E. Pnevmatikakis, P. E. Latham, J. P. Cunningham, and A. K. Churchland. Excitatory and inhibitory subnetworks are equally selective during decision-making and emerge simultaneously during learning. Neuron, 105(1): 165–179, 2020.

G. K. Ocker, Y. Hu, M. A. Buice, B. Doiron, K. Josić, R. Rosenbaum, and E. Shea-Brown. From the statistics of connectivity to the statistics of spike times in neuronal networks. Current opinion in neurobiology, 46:109–119, 2017.

H. Ozeki, I. M. Finn, E. S. Schaffer, K. D. Miller, and D. Ferster. Inhibitory stabilization of the cortical network underlies visual surround suppression. Neuron, 62 (4):578–592, 2009.

A. M. Packer and R. Yuste. Dense, unspecific connectivity of neocortical parvalbumin-positive interneurons: a canonical microcircuit for inhibition? Journal of Neuroscience, 31(37):13260–13271, 2011.

V. Pernice, B. Staude, S. Cardanobile, and S. Rotter. How structure determines correlations in neuronal networks. PLoS computational biology, 7(5):e1002059, 2011.

A. Renart, N. Brunel, and X.-J. Wang. Mean-field theory of irregularly spiking neuronal populations and working memory in recurrent cortical networks. Computational neuroscience: A comprehensive approach, pages 431–490, 2004.

A. Renart, J. De La Rocha, P. Bartho, L. Hollender, N. Parga, A. Reyes, and K. D. Harris. The asynchronous state in cortical circuits. science, 327(5965):587–590, 2010.

R. Rosenbaum, M. A. Smith, A. Kohn, J. E. Rubin, and B. Doiron. The spatial structure of correlated neuronal variability. Nature neuroscience, 20(1):107–114, 2017.

Y. Roudi, B. Dunn, and J. Hertz. Multi-neuronal activity and functional connectivity in cell assemblies. Current opinion in neurobiology, 32:38–44, 2015.

D. B. Rubin, S. D. Van Hooser, and K. D. Miller. The stabilized supralinear network: a unifying circuit motif underlying multi-input integration in sensory cortex. Neuron, 85(2):402–417, 2015.

S. Sadeh and C. Clopath. Inhibitory stabilization and cortical computation. Nature Reviews Neuroscience, 22(1):21–37, 2021.

A. Sanzeni, B. Akitake, H. C. Goldbach, C. E. Leedy, N. Brunel, and M. H. Histed. Inhibition stabilization is a widespread property of cortical networks. Elife, 9:e54875, 2020.

A. C. Schwarze and M. A. Porter. Motifs for processes on networks. SIAM Journal on Applied Dynamical Systems, 20(4):2516–2557, 2021.

R. Tremblay, S. Lee, and B. Rudy. Gabaergic interneurons in the neocortex: from cellular properties to circuits. Neuron, 91(2):260–292, 2016.

J. Trousdale, Y. Hu, E. Shea-Brown, and K. Josić. Impact of network structure and cellular response on spike time correlations. PLoS computational biology, 8(3): e1002408, 2012.

M. V. Tsodyks, W. E. Skaggs, T. J. Sejnowski, and B. L. McNaughton. Paradoxical effects of external modulation of inhibitory interneurons. Journal of neuroscience, 17 (11):4382–4388, 1997.

G. G. Turrigiano. The self-tuning neuron: synaptic scaling of excitatory synapses. Cell, 135(3):422–435, 2008.

A. E. Urai, B. Doiron, A. M. Leifer, and A. K. Churchland. Large-scale neural recordings call for new insights to link brain and behavior. Nature neuroscience, 25 (1):11–19, 2022.

S. Wiggins. Introduction to applied nonlinear dynamical systems and chaos, volume 2. Springer, 2003.

P. Znamenskiy, M.-H. Kim, D. R. Muir, M. F. Iacaruso, S. B. Hofer, and T. D. Mrsic-Flogel. Functional selectivity and specific connectivity of inhibitory neurons in primary visual cortex. Biorxiv, page 294835, 2018.

